# Soil invertebrate biomass estimation from image-derived body area and contour length

**DOI:** 10.64898/2026.10.07.757327

**Authors:** Ajuan Zhang, Jing-Zhong Lu, Andrey G. Zuev, Bingxue Zhu, Nicole Scheunemann, Cornelia Wiesener, Andreas Kauk, Birgit Balkenhol, Julia Noe, Sebastian Moll, Simin Li, Anton Potapov, Clément Schneider

**Affiliations:** Senckenberg Museum of Natural History Görlitz, Department of Soil Zoology, Am Museum 1, 02826 Görlitz, Germany; German Centre for Integrative Biodiversity Research (iDiv) Halle-Jena-Leipzig, Puschstraße 4, 04103 Leipzig, Germany; State Key Laboratory of Black Soils Conservation and Utilization, Northeast Institute of Geography and Agroecology, CAS, Changchun 130102, China; Research Center of Solid Waste Pollution and Prevention, Nanjing Institute of Environmental Science, Ministry of Ecology and Environment, Nanjing 210042, PR China; International Institute Zittau, TUD Dresden University of Technology, 02763 Zittau, Germany

**Keywords:** body area, contour length, dry mass, ecosystem function, fresh mass, macrofauna, mesofauna, soil invertebrate biomass

## Abstract

1. Soil invertebrate biomass is closely linked to ecosystem processes, yet its routine quantification remains constrained by the effort required to weigh large numbers of small individuals. Length–mass relationships provide an indirect alternative, but body length can be difficult to standardise and extract automatically from images of curved, coiled or contracted organisms. Projected body area may provide a more tractable image-derived predictor, but its performance across diverse soil-faunal taxa and at the community scale remains insufficiently evaluated.
2. Here, we developed a framework using projected body area and contour length to estimate fresh and dry mass from measurements of 1,792 soil invertebrates representing 15 higher-level taxa. We fitted taxon-specific models for six well-represented groups and general models across taxa, and evaluated predictive performance using leave-one-out cross-validation. We then assessed cross-dataset transferability by comparing our data with those of Lemoine et al. (2026) and pooled the two datasets to derive combined models. Finally, we tested whether individual predictions could be scaled to independently measured community biomass.
3. Taxon-specific models generally reduced prediction error relative to general models, with mean absolute percentage error (MAPE) values of 6.9–37.3% for fresh mass and 7.6–30.2% for dry mass. General models retained MAPEs of 27.2% and 22.4%, respectively. Comparisons of our dataset with that of Lemoine et al. (2026) showed no detectable differences in six of eight taxon-by-mass slopes. Although intercepts frequently differed, no overall dataset effect was detected after accounting for taxa. Pooling both datasets broadened model development, while taxon-specific models generally retained lower error than the combined general model. Finally, community-level predictions tracked measured biomass across all model sets (*R*^2^ = 0.87– 0.99), but absolute error varied among models and faunal groups (MAPE = 9.6–52.5%).
4. Body area and contour length extracted from images therefore provide a practical basis for scaling soil invertebrate biomass estimation from individuals to communities. Integration with automated imaging and segmentation tools such as *flatbug*, BIODISCOVER and CollembolAI could extend soil invertebrate monitoring at larger scales beyond abundance and taxonomic composition to biomass, while providing scalable inputs for analyses of food webs and general ecosystem ecology.

## 1 | INTRODUCTION

Soil invertebrates are integral to terrestrial ecosystems, contributing to organic matter processing, nutrient cycling and other biogeochemical processes that underpin ecosystem functioning (Crowther *et al*. 2019; Eisenhauer *et al*. 2026; Frouz 2018; Wu *et al*. 2025). Yet assessments of soil invertebrate communities have traditionally focused on taxonomic diversity and abundance, with biomass receiving comparatively less attention (Ellwood & Foster 2004; Winfree *et al*. 2025). Because body mass influences individual energy requirements, biomass provides functional information beyond diversity and abundance and is essential for estimating energy fluxes in soil food webs (Bonfanti *et al*. 2024; Brown *et al*. 2004; Gauzens *et al*. 2019; Jochum *et al*. 2021). Experimental reductions in invertebrate biomass can alter ecosystem processes and multifunctionality independently of species loss (Eisenhauer *et al*. 2023), while declines in terrestrial arthropod abundance and biomass have been documented across ecosystems (van Klink *et al*. 2020; Seibold *et al*. 2019). Accurate and scalable estimates of soil invertebrate biomass are therefore increasingly important for community assessment and process-based analyses of food webs and ecosystem functioning under environmental change.

Direct measurement of invertebrate biomass is labour-intensive, particularly for small-bodied taxa, because large numbers of specimens must be sorted, identified and weighed individually (Ärje *et al*. 2020; Coulis & Joly 2017; Penell *et al*. 2018). This hampers biomass assessments in biodiversity surveys, replicated experiments and long-term monitoring. Consequently, individual body mass is often estimated indirectly from morphological traits, most commonly through taxon-specific length– mass relationships, and subsequently combined with abundance to derive community biomass (Sohlström *et al*. 2018). However, two limitations constrain the broader application of this approach. First, mass–size relationships vary among taxa, body forms and geographic regions, such that predictive accuracy can decline when regression parameters are transferred beyond the groups or conditions for which they were calibrated (Höfer & Ott 2009; Martin *et al*. 2014; Schoener 1980). Second, linear body length can be difficult to standardise for soil invertebrates that adopt curved, coiled or contracted postures, particularly after handling or preservation, making measurements more time-consuming and potentially less reproducible and harder to automate (Coulis & Joly 2017). These limitations restrict the efficiency with which conventional morphometric approaches can be applied to large and morphologically diverse soil-faunal datasets.

Recent advances in automated imaging and computer vision offer an opportunity to overcome some of these constraints. Image-based systems can now detect, count and classify preserved soil invertebrates with high accuracy, reducing reliance on manual processing and expert-based identification (Ärje *et al*. 2020; Sys *et al*. 2022). Importantly, the same images can provide standardised morphometric information for each individual, including projected body area and contour-based measurements. Unlike a single linear dimension, projected area integrates variation in both body length and width, while contour metrics can provide complementary information on body shape. Previous studies have shown that two-dimensional area and related image-derived size descriptors can predict invertebrate mass in several taxa (Ärje *et al*. 2020; Coulis & Joly 2017; Lemoine *et al*. 2026), although predictive performance can vary substantially among species and body-size classes. These developments suggest that routine digital images could provide a practical basis for estimating individual biomass while reducing dependence on separate manual morphometric measurements.

Despite this progress, it remains unclear how consistently image-derived morphology predicts biomass across soil invertebrates that differ widely in body size and form, whether taxon-specific calibration improves prediction relative to a general cross-taxon model, and whether individual-level prediction errors remain acceptable when scaled to community biomass. Furthermore, no area-mass regressions are currently available for small-sized but very abundant soil invertebrates such as springtails and mites, which account for 95% of soil arthropods globally (Rosenberg *et al*. 2023).

Here, we developed an image-based framework for estimating fresh and dry mass from projected body area and contour length (Figure 1). Specifically, we (1) fitted taxon-specific models for six major meso-and macrofaunal groups (i.e., Lumbricina, Araneae, Oribatida, Myriapoda, Collembola, Formicidae) and general models across taxa; (2) evaluated cross-dataset transferability by comparing and pooling our data with those of Lemoine et al. (2026); and (3) independently tested whether individual mass predictions could be scaled to community biomass. This framework links image-derived morphology to biomass from individuals to communities and provides a basis for integration with increasingly automated soil-fauna imaging workflows.

**FIGURE 1.**
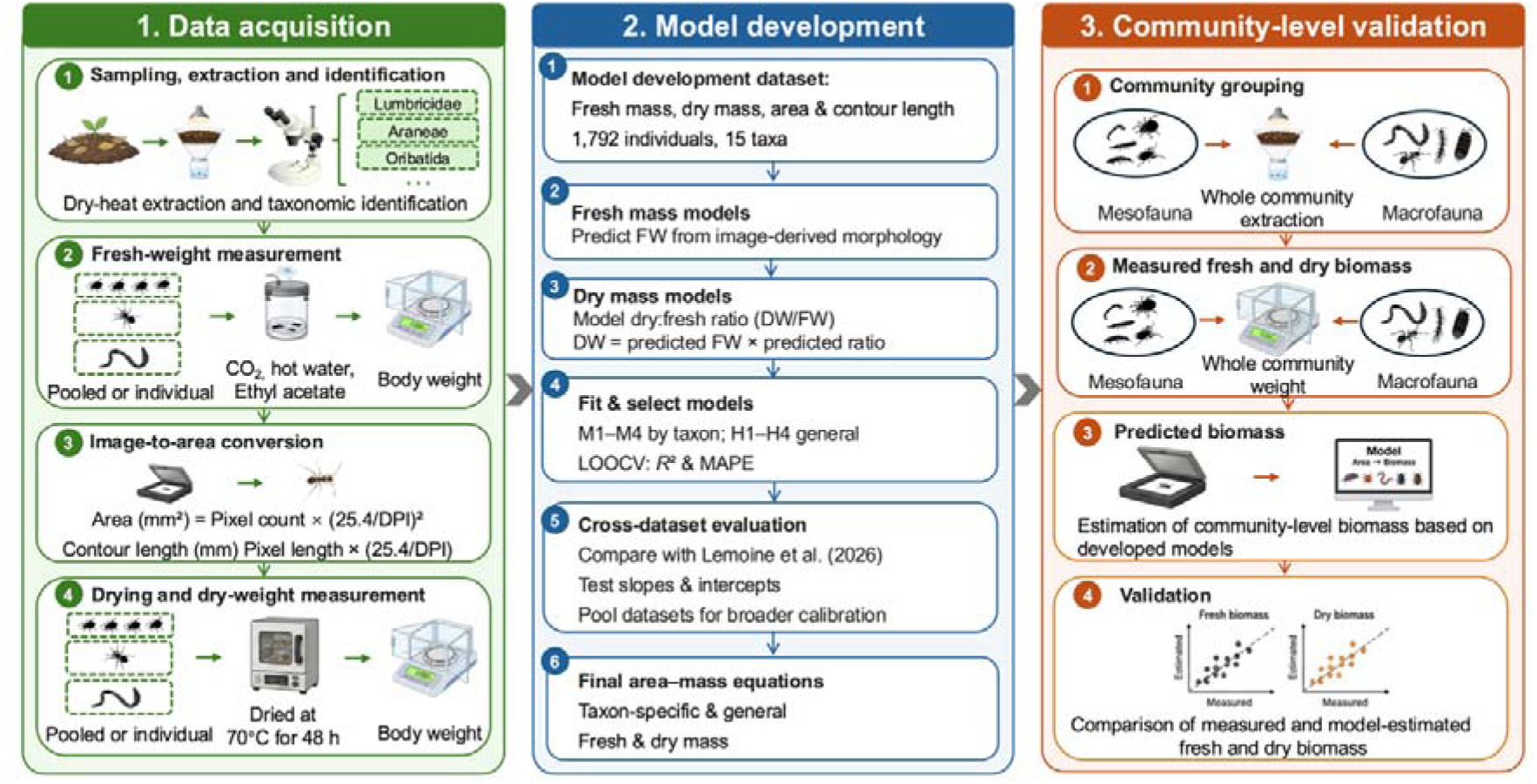
Workflow for image-based estimation and validation of soil invertebrate biomass. The workflow comprised three stages. (1) **Data acquisition:** soil invertebrates were sampled, identified and weighed for fresh mass, either individually or in pooled groups of similarly sized individuals. Specimens were then imaged to quantify projected body area and contour length, dried and reweighed for dry mass. (2) **Model development:** taxon-specific and general models were fitted for fresh mass and the dry:fresh mass ratio and evaluated using leave-one-out cross-validation. Cross-dataset transferability was further assessed using data from Lemoine et al. (2026), and both datasets were pooled to broaden calibration. (3) **Community-level validation:** the resulting equations were applied to independent mesofauna and macrofauna communities, and predicted fresh and dry biomass were compared with directly measured community biomass. Soil animal icons were created by Svenja Meyer.

## 2 | MATERIALS AND METHODS

### 2.1 Sampling and fresh body mass measurements

#### Sampling

For model development, 1,792 soil invertebrates were obtained between May and September 2024 in the surroundings and collections of Görlitz, Saxony, Germany. Most taxa were collected directly by hand or extracted from field-collected litter using dry-heat Berlese funnels for 7 days. Collembola were obtained from laboratory cultures maintained at Senckenberg Görlitz. All specimens were alive before fresh mass measurement and were held for no longer than 24 h before weighing.

#### Fresh body mass measurements

To minimise movement during weighing and obtain stable fresh-mass measurements, specimens were immobilised immediately before weighing using taxon-specific procedures. Araneae and Oribatida were exposed to ethyl acetate vapour, Formicidae to CO_2_, and Collembola to either CO_2_ or ethyl acetate vapour. Enchytraeidae, Dermaptera and Diptera were treated briefly with hot water, whereas Hemiptera, Coleoptera and other arthropods (Opiliones, Prostigmata and Coleoptera larvae) were treated with either ethyl acetate vapour or hot water. Lumbricina, Isopoda and Myriapoda were sufficiently inactive for stable measurement and were therefore weighed alive without prior immobilisation. Most sufficiently large specimens were weighed individually. Collembola and Oribatida, however, were generally below the reliable range for individual weighing and were weighed collectively in groups of similarly sized individuals. Group fresh mass was divided by the number of individuals to obtain mean individual fresh mass, with each group mean treated as one observation in subsequent analyses. Because

Collembola were particularly difficult to handle owing to their minute size and rapid fresh-mass loss after transfer, two size-matched sets containing equal numbers of individuals were prepared for each species: one for fresh-and subsequent dry-mass measurement and the other for imaging. Specimens designated for imaging were preserved in 70% ethanol until scanning.

### 2.2 Image acquisition, area extraction and dry body mass measurement

#### Image acquisition

Following the Soil BON Foodweb imaging protocol (Potapov *et al*. 2022), individuals from each sample or taxon were distributed on Petri dishes for subsequent image-based body-size estimation. Taxonomic groups were scanned separately in 70% ethanol in 9-cm plastic Petri dishes against a white background using an EPSON Perfection V600 flatbed scanner connected to a PC. All scans were acquired at 2400 dpi under standardised professional-mode settings (reflective document type, document table source, document auto exposure, 48-bit colour, JPEG export, and unsharp mask enabled).

#### Image processing and area calculation

Binary masks were manually generated from the scanned community images using Krita (Version: 5.3.2; Krita Foundation, https://www.kde.org/), a raster image editor. Images were first desaturated and converted to binary images, with objects represented as black pixels against a white background. Where adjacent objects remained connected after thresholding, they were manually separated using a 2-pixel-wide brush. Small specks, dust and other debris visible in the original images were manually erased during this procedure. Individual objects, including animals and non-animal particles, were subsequently detected automatically using Python and the OpenCV library (v4.12.0, Open Source Vision Foundation, https://opencv.org/). For each separated object, OpenCV extracted its bounding box and outline in pixel coordinates. The Python code used for automated object detection and morphometric extraction is available from figshare (Code availability). Object area was quantified as the number of black pixels within the corresponding binary mask, whereas contour length was estimated from the number of horizontal and vertical transitions between object and background pixels along the object boundary. Pixel-based measurements were converted to physical units according to the scan resolution of 2400 dpi. Given that 1 inch = 25.4 mm, the linear dimension of one pixel corresponded to 25.4/2400 = 0.0105833 mm. Object area and contour length were therefore calculated as (See Supplementary Figures S1–S3 for the image-processing workflow, pixel-based calculation of projected body area and contour length, and binary-mask preparation in Krita):

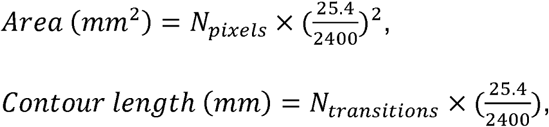

where *N*_pixels_ is the number of black pixels assigned to an object and *N*_transitions_ is the number of horizontal and vertical object–background transitions along its boundary.

Each detected animal was then manually assigned to its corresponding taxonomic group. Non-animal objects were classified separately as either larger dust/debris particles or small indistinct (‘fuzzy’) particles.

#### Dry body mass measurements

Specimens used for dry mass determination were oven-dried at 70 °C for 48 h and reweighed.

### 2.3 Model development and selection

#### Model development data

Each model-development observation comprised projected body area (*A*, mm^2^) and contour length (*L*, mm, measured as the contour length of the digitised body outline), fresh mass (FW, μg) and dry mass (DW, μg). Taxon-specific models were fitted for six higher-level groups with sufficient replication: Lumbricina, Araneae, Oribatida, Myriapoda, Collembola and Formicidae. A separate General dataset contained 551 observations representing 15 taxa and was analysed using hierarchical models. All predictor and response variables were required to be positive before natural-log transformation.

#### Allometric model formulation

Relationships between organismal size traits are commonly described by allometric power laws (Huxley *et al*. 1993), and this framework has been widely applied to estimate invertebrate body mass from morphometric measurements (Coulis & Joly 2017; Hódar 1996):

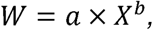

where W is body mass, *X* is a morphometric predictor (Area or Length), *a* is a normalisation constant, and *b* is the scaling exponent. A natural-log (ln) transformation linearises this relationship,

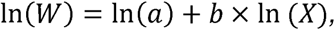

which additionally stabilises the variance of residuals, a property of central importance for body-mass data, in which absolute measurement and biological variance typically increase with body size.

We initially considered 12 candidate models: the four ln–ln allometric models retained in the final analysis and eight additional linear or quadratic models fitted to untransformed Area and Length. The eight raw-scale models were excluded for both conceptual and empirical reasons. Unlike the ln-transformed power-law models, they were purely empirical and lacked a biologically meaningful growth form. In particular, raw-scale quadratic models can imply unrealistic changes in mass beyond the observed size range, while both linear and quadratic fits can generate biologically impossible negative predictions. In preliminary comparisons, none of these models outperformed their ln–ln counterparts, supporting their exclusion from the final candidate set.

#### Taxon-specific models

Four candidate models were considered as reduced forms of a second-order polynomial in ln(*A*) and ln(*L*):

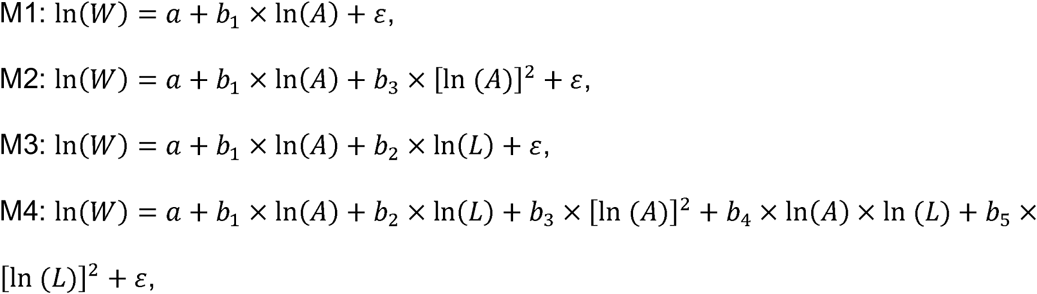

The four candidate models represented reduced forms of M4, with M1–M3 retaining different subsets of its terms.

#### General models across taxa

For the cross-taxon “general” model, rows from all taxa were not pooled into an ordinary regression, because the intercept and/or allometric exponent linking body size to mass are expected to differ systematically among taxa (e.g., a sclerotised exoskeleton in Coleoptera vs. the soft cuticle of Collembola). Four hierarchical models (H1–H4) were therefore defined by varying two structural components: whether contour length was included as an additional fixed-effect predictor and whether the effect of ln(*A*) was allowed to vary among taxa.

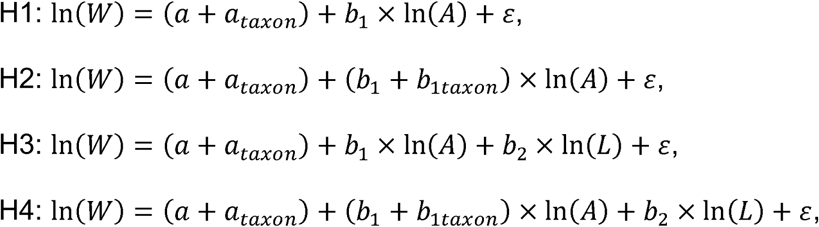

where *a*, *b*_1_ and *b*_2_ are fixed effects shared across taxa, and *a*_taxon_ and *b*_₁taxon_ are taxon-specific random deviations in intercept and area slope. H1 and H3 therefore allowed taxa to differ in intercept, whereas H2 and H4 additionally allowed the area-mass slope to vary among taxa. Random slopes were restricted to ln(*A*) to limit model complexity and because ln(*A*) and ln(*L*) were strongly correlated.

#### Estimating dry mass from the dry:fresh mass ratio

Rather than modelling DW independently, we modelled the dry:fresh mass ratio (r = DW/FW), which is bounded between 0 and 1. Ratios ≤0 or ≥1 were considered invalid, although no observations met this criterion. The ratio was modelled on the logit scale, logit(*r*) = ln[*r*/(1 − *r*)], using the same taxon-specific and hierarchical candidate structures described above.

Predictions were transformed back using the inverse-logit function, and dry mass was reconstructed as DW = FW × *r*. This formulation constrains predicted dry mass not to exceed predicted fresh mass.

#### Model evaluation and selection

All taxon-specific and hierarchical candidate models were evaluated using leave-one-out cross-validation (LOOCV). Each model-development observation was omitted in turn, the model was refitted to the remaining observations, and mass was predicted for the omitted observation on the original mass scale. Predictive performance was quantified using LOOCV *R*^2^ and mean absolute percentage error (MAPE). As supplementary diagnostics, MAPE was also calculated separately for the smaller and larger 50% of individuals to assess size-dependent prediction error, and aggregate percentage error was calculated from the summed predicted and observed masses to evaluate performance at the community-biomass level.

MAPE was used as the primary model-selection criterion because it directly measures relative prediction error. For each taxon, the model with the lowest MAPE was identified as the best predictive model. To avoid retaining unnecessary complexity, the parsimonious model was defined as the model with the fewest estimated parameters among those with MAPE within 1 percentage point of the minimum. The parsimonious model was used in the main analyses, while complete results for both best predictive and parsimonious models are reported in Tables S1 and S2.

### 2.4 Evaluation of model generality using an independent dataset

To assess the generality of our area–mass relationships, we compared our data with an independently collected, methodologically similar dataset (Lemoine et al., 2026; n = 1,153 specimens; CIRAD Dataverse, https://doi.org/10.18167/DVN1/SCH4AV). Both studies quantified body area from scanned images and included four shared groups: Araneae, Hymenoptera (represented by Formicidae in our study), Myriapoda (i.e., Diplopoda and Chilipoda) and Lumbricina (represented by Megascolecidae, Glossoscolecidae, Rhinodrilidae, Eudrilidae and Octochaetidae in the Lemoine et al. dataset). Mass units were standardised from mg to µg and larval stages were excluded. For direct comparability, cross-dataset analyses used an area-only ln-linear model common to both studies, corresponding to our M1 model (i.e., ln(W) = *a* + *b*₁ln(*A*) + ε). For each shared group and mass type (fresh and dry), we compared three nested models: a pooled model with common slope and intercept, a model with dataset-specific intercepts and a common slope, and a model with dataset-specific slopes and intercepts. Nested-model *F*-tests assessed differences in slope and, assuming a common slope, intercept between datasets. At the multi-taxon level, we pooled all taxa from both datasets and fitted linear mixed-effects models with taxon-specific random intercepts and slopes. Models with and without data source as a fixed effect were compared using a likelihood-ratio test to assess systematic differences between datasets after accounting for taxon-level variation. Combined taxon-specific and general M1 models were fitted to pooled datasets for fresh and dry mass and evaluated by LOOCV.

### 2.5 Independent validation at the community scale

An independent set of soil-invertebrate communities was collected from litter near Görlitz in autumn 2026. Mesofauna were extracted using dry-heat Berlese funnels, whereas macrofauna, including earthworms, were handpicked. Mesofauna comprised Collembola, Oribatida, Mesostigmata, Prostigmata, Astigmata, Diplura and Symphyla, whereas macrofauna comprised Chilopoda, Araneae, Isopoda, Diplopoda, Lumbricina, Diptera, Coleoptera, Lepidoptera, Formicidae, Sternorrhyncha, Heteroptera and Thysanoptera. Fresh biomass was measured for each mesofauna and macrofauna community as a whole, then imaged following the same protocol as the model-development dataset, after which samples were oven-dried and reweighed. Directly measured fresh and dry biomass was therefore available for 10 mesofauna and 10 macrofauna communities. We then applied the same manual thresholding method and object splitting to create the binary masks. Residual dust and soil particles visible in the images were retained despite efforts to minimise their presence. Individual objects were automatically detected using

OpenCV, and their bounding boxes and segmentation coordinates were stored in JSON files following the COCO annotation format. Each object was then cropped into a separate image, with its object ID retained in the filename. Animals and non-animal particles were manually classified by placing the cropped images into label-specific folders, and the resulting labels were mapped back to the COCO dataset. Non-animal particles were excluded from biomass prediction and were not included when individual predictions were summed to estimate community biomass.

To test whether individual mass predictions could be reliably scaled to community biomass, we applied taxon-specific equations where available and the general model otherwise. Predicted individual masses were summed within each validation sample and compared with independently measured fresh and dry community biomass.

Validation was conducted separately for macrofauna and mesofauna because mesofauna comprised many small-bodied individuals for which individual mass predictions were less precise. Agreement between predicted and measured biomass was evaluated using *R*^2^, MAPE and the mean predicted-to-measured biomass ratio, with the latter used to assess systematic over-or underestimation. For comparison, validation was repeated separately using the General model and the combined-dataset equations.

## 3 RESULTS

### 3.1 Taxonomic coverage and body-size range

The dataset comprised 1,792 soil invertebrates assigned to 15 taxonomic groups (Table 1). Sample sizes varied considerably among groups, with Collembola having the largest sample size, followed by Oribatida, Myriapoda, Isopoda and Formicidae.

**Table 1.**
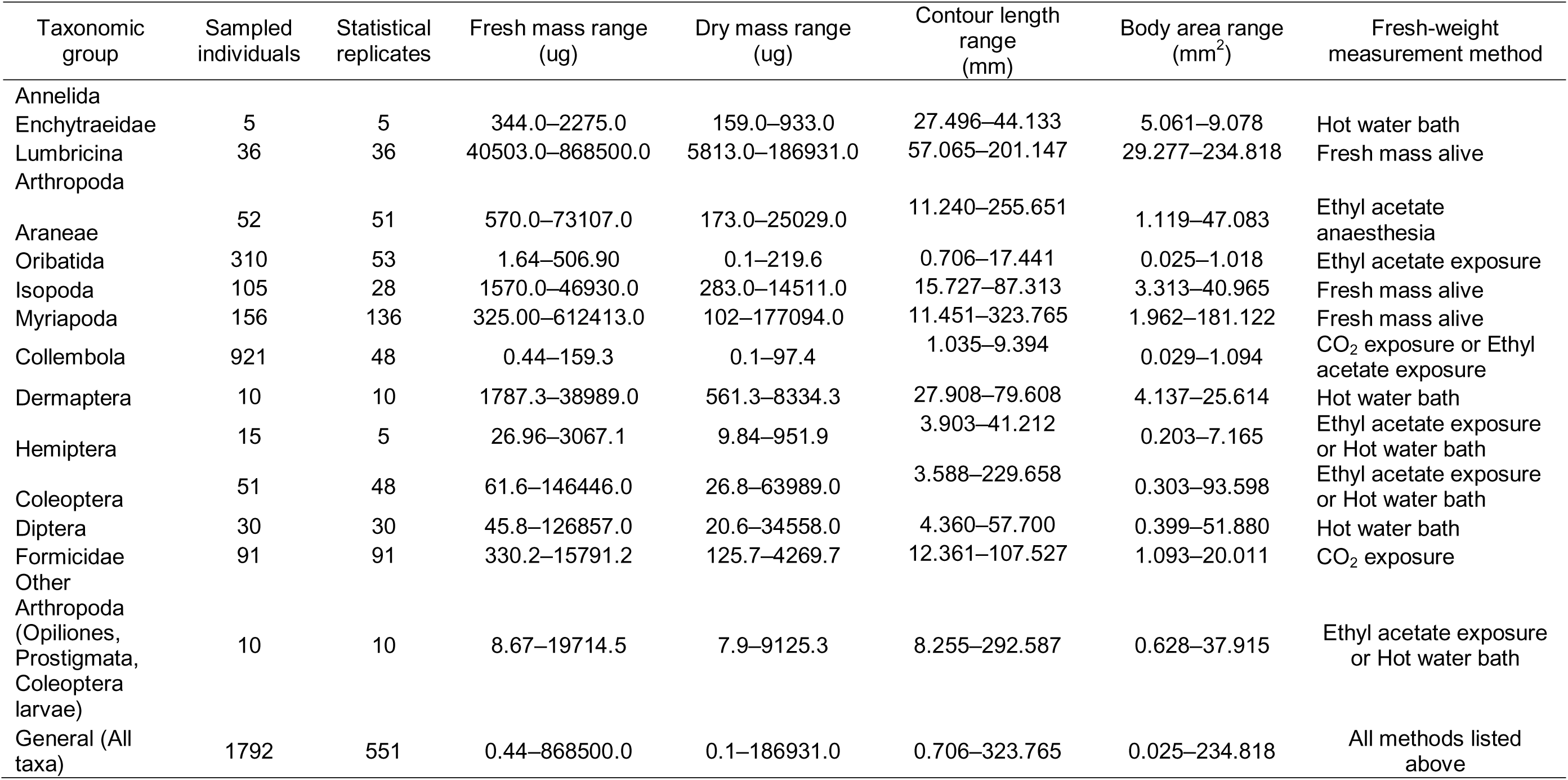
Summary of soil invertebrates used to develop body area–biomass regression models. For each taxonomic group, the table reports the number of individuals sampled, the number of statistical replicates used for analysis, the observed ranges of fresh mass, dry mass and body area, and the method used for fresh-weight measurement. Weight ranges are given in µg and body area in mm^2^. Statistical replicates refer to independent weight–area observations used for model fitting; for small-bodied taxa, a replicate represents pooled individuals of similar size that were weighed together and converted to an individual-level estimate. Fresh-weight measurement method denotes the handling, anaesthesia or killing procedure applied immediately before fresh-weight determination.

Enchytraeidae, Hemiptera, Dermaptera and the pooled Other Arthropoda group (Opiliones, Prostigmata and Coleoptera larvae) were represented by comparatively few individuals.

Measured body masses spanned approximately six orders of magnitude across taxa. Fresh mass ranged from <1 µg in Collembola to 868,500 µg in Lumbricina, whereas dry mass ranged from 0.1 to 186,931 µg, with the maximum again recorded in Lumbricina (Table 1). Contour length ranged from <1 mm in Oribatida to >300 mm in Myriapoda, while projected body area ranged from 0.025 mm^2^ in Oribatida to 234.8 mm^2^ in Lumbricina (Table 1).

### 3.2 Regressions

Fresh-mass models differed substantially in predictive performance among the six taxa (Table 2, Figures 2 and S4). Lumbricina had the lowest prediction error (M3; LOOCV *R*^2^ = 0.95, MAPE = 6.9%), followed by Formicidae (M3; *R^2^* = 0.95, MAPE = 16.4%). Araneae, Myriapoda and Oribatida showed similar MAPEs (23.2–23.9%; M3 for all three), although their *R^2^* values differed (0.93, 0.94 and 0.85, respectively).

**FIGURE 2.**
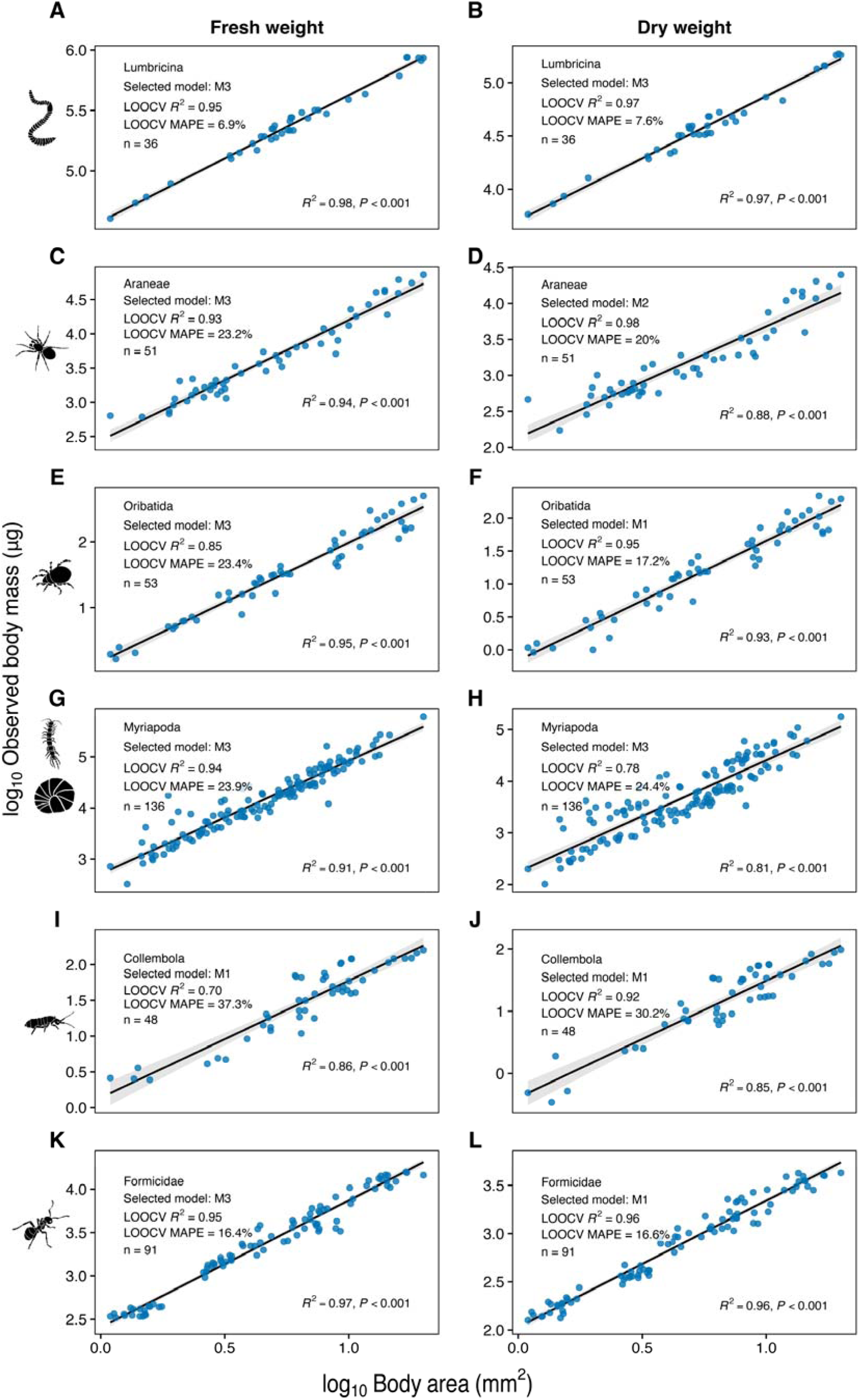
Taxon-specific relationships between image-derived body area and fresh and dry mass of soil invertebrates. Fresh mass (left) and dry mass (right) are shown for Lumbricina (**A, B**), Araneae (**C, D**), Oribatida (**E, F**), Myriapoda (**G, H**), Collembola (**I, J**) and Formicidae (**K, L**). Points represent statistical observations; for Oribatida and Collembola, these correspond to group-derived mean masses from pooled individuals. Black lines and grey bands show simple log_10_–log_10_ regressions between body area and observed mass with 95% confidence intervals. Upper-left annotations report the selected taxon-specific model and its leave-one-out cross-validation (LOOCV) performance, whereas lower-right annotations report the *R*^2^ and *P* value of the simple area–mass regression shown in each panel. Soil animal icons were created by S. Meyer.

**Table 2.** Selected parsimonious models for estimating fresh mass (FW, µg) of soil invertebrates from image-derived body area (*A*, mm^2^) and contour length (*L*, mm). M1–M4 and H1–H4 are defined in the Methods. Selected models were the simplest candidates within 1 percentage point of the minimum LOOCV MAPE. En dashes indicate excluded terms. LOOCV *R*^2^ and MAPE were calculated on the original mass scale after back-transformation.

| Taxon/group | Model | Intercept<br>$a$ | $\ln(A)$<br>$b_1$ | $\ln(L)$<br>$b_2$ | $[\ln(A)]^2$<br>$b_3$ | $\ln(A) \times \ln(L)$ $b_4$ | $[\ln(L)]^2$<br>$b_5$ | LOOCV $R^2$ | LOOCV<br>MAPE (%) |
| --- | --- | --- | --- | --- | --- | --- | --- | --- | --- |
| Lumbricina | M3 | 6.5639 | 1.6570 | -0.3715 | – | – | – | 0.950 | 6.9% |
| Araneae | M3 | 7.3900 | 1.7162 | -0.6139 | – | – | – | 0.933 | 23.2% |
| Oribatida | M3 | 7.0722 | 1.8398 | -0.5926 | – | – | – | 0.845 | 23.4% |
| Myriapoda | M3 | 7.3443 | 1.8916 | -0.7106 | – | – | – | 0.935 | 23.9% |
| Collembola | M1 | 5.0991 | 1.3073 | – | – | – | – | 0.704 | 37.3% |
| Formicidae | M3 | 6.6616 | 1.7210 | -0.4219 | – | – | – | 0.953 | 16.4% |
| General (All taxa) | H4 | 6.8998 | 1.7232 | -0.5836 | – | – | – | 0.970 | 27.2% |
| Lumbricina: FW = $\exp[6.5639 + 1.6570 \times \ln(A) - 0.3715 \times \ln(L)]$ | | | | | | | | | |
| Araneae: FW = $\exp[7.3900 + 1.7162 \times \ln(A) - 0.6139 \times \ln(L)]$ | | | | | | | | | |
| Oribatida: FW = $\exp[7.0722 + 1.8398 \times \ln(A) - 0.5926 \times \ln(L)]$ | | | | | | | | | |
| Myriapoda: FW = $\exp[7.3443 + 1.8916 \times \ln(A) - 0.7106 \times \ln(L)]$ | | | | | | | | | |
| Collembola: FW = $\exp[5.0991 + 1.3073 \times \ln(A)]$ | | | | | | | | | |
| Formicidae: FW = $\exp[6.6616 + 1.7210 \times \ln(A) - 0.4219 \times \ln(L)]$ | | | | | | | | | |
| General: FW = $\exp[6.8998 + 1.7232 \times \ln(A) - 0.5836 \times \ln(L)]$ | | | | | | | | | |

Collembola showed the weakest predictive performance (M1; *R^2^* = 0.70, MAPE = 37.3%). The general model (H4) achieved a high LOOCV *R^2^* of 0.97 but a MAPE of 27.2%, exceeding the prediction error of all taxon-specific models except Collembola (Table 2, Figures 2 and 3).

**FIGURE 3.**
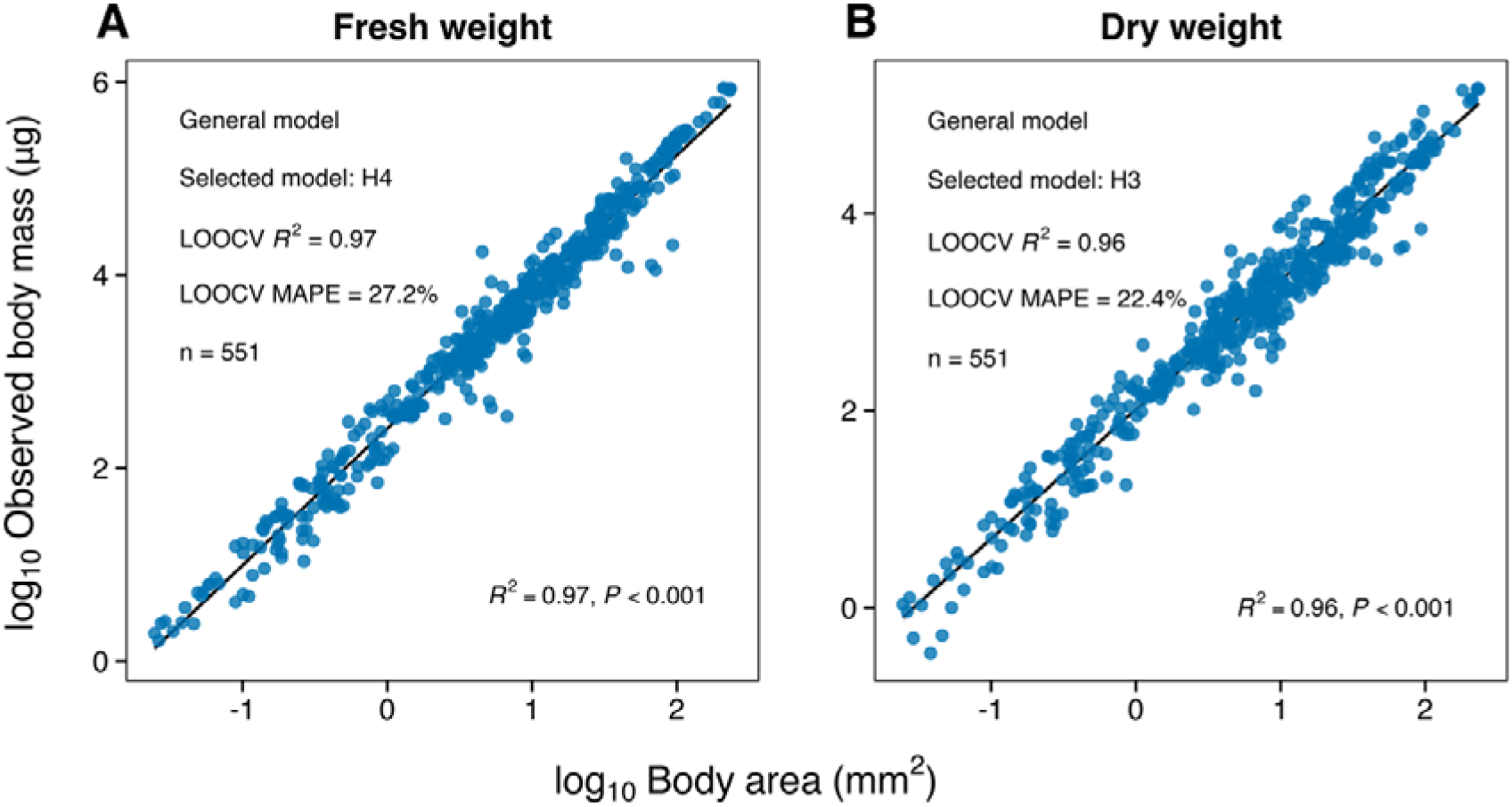
Relationships between image-derived body area and fresh and dry mass in the general dataset. Fresh mass (A) and dry mass (B) are shown across all taxa included in the general model. Points represent statistical observations. Black lines and grey bands show simple log_10_–log_10_ regressions between body area and observed mass with 95% confidence intervals. Upper-left annotations report the selected hierarchical model and its leave-one-out cross-validation (LOOCV) performance, whereas lower-right annotations give the *R*^2^ and *P* value of the simple regression shown for visualisation.

Dry-mass models also varied substantially in predictive performance among taxa (Table 3, Figures 2 and S5). LOOCV *R*^2^ was higher for dry than for fresh mass in five of the six taxa, whereas MAPE showed no consistent improvement. Lumbricina had the lowest relative prediction error (M3; *R*^2^ = 0.97, MAPE = 7.6%), followed by Formicidae (M1; *R*^2^ = 0.96, MAPE = 16.6%), Oribatida (M1; *R*^2^ = 0.95, MAPE = 17.2%) and Araneae (M2; *R*^2^ = 0.98, MAPE = 20.0%). Myriapoda and Collembola had MAPEs of 24.4% and 30.2% (Myriapoda *R*^2^ = 0.78, Collembola *R^2^* = 0.92). The general model (H3) achieved an *R*^2^ of 0.96 and a MAPE of 22.4%, with higher relative prediction error than four of the six taxon-specific models (Table 3, Figures 2, 3 and S6).

**Table 3.** Selected parsimonious models for estimating dry mass (DW, µg) of soil invertebrates from image-derived body area (*A*, mm^2^) and contour length (*L*, mm), expressed as the dry:fresh mass ratio. The dry:fresh mass ratio (DW/FW) was modelled on the logit scale, and dry mass was reconstructed as predicted FW × plogis(linear predictor), ensuring DW ≤ FW. Candidate models M1– M4 and hierarchical models H1–H4 are defined in the Methods and followed the same selection rule as in Table 2. En dashes indicate terms not included in the selected model. LOOCV *R*^2^ and MAPE were calculated on the reconstructed dry-mass scale.

| Taxon/group | Model | Intercept<br>$a$ | $\ln(A)$<br>$b_1$ | $\ln(L)$<br>$b_2$ | $[\ln(A)]^2$<br>$b_3$ | $\ln(A) \times \ln(L)$<br>$b_4$ | $[\ln(L)]^2$<br>$b_5$ | LOOCV $R^2$ | LOOCV<br>MAPE (%) |
| --- | --- | --- | --- | --- | --- | --- | --- | --- | --- |
| Lumbricina | M3 | -1.5360 | 0.4460 | -0.4485 | – | – | – | 0.969 | 7.6% |
| Araneae | M2 | 0.6910 | -1.071 | – | 0.1827 | – | – | 0.979 | 20.0% |
| Oribatida | M1 | -0.1222 | -0.0071 | – | – | – | – | 0.953 | 17.2% |
| Myriapoda | M3 | 3.5569 | 0.8500 | -1.4728 | – | – | – | 0.783 | 24.4% |
| Collembola | M1 | 0.4085 | 0.2965 | – | – | – | – | 0.923 | 30.2% |
| Formicidae | M1 | -0.3081 | -0.2366 | – | – | – | – | 0.959 | 16.6% |
| General (All taxa) | H3 | 1.4023 | 0.4203 | -0.8017 | – | – | – | 0.961 | 22.4% |
| Lumbricina: $\text{DW} = \text{FW} \times \text{plogis}[-1.5360 + 0.4460 \times \ln(A) - 0.4485 \times \ln(L)]$ | | | | | | | | | |
| Araneae: $\text{DW} = \text{FW} \times \text{plogis}[0.6910 - 1.0710 \times \ln(A) + 0.1827 \times [\ln(A)]^2]$ | | | | | | | | | |
| Oribatida: $\text{DW} = \text{FW} \times \text{plogis}[-0.1222 - 0.0071 \times \ln(A)]$ | | | | | | | | | |
| Myriapoda: $\text{DW} = \text{FW} \times \text{plogis}[3.5569 + 0.8500 \times \ln(A) - 1.4728 \times \ln(L)]$ | | | | | | | | | |
| Collembola: $\text{DW} = \text{FW} \times \text{plogis}[0.4085 + 0.2965 \times \ln(A)]$ | | | | | | | | | |
| Formicidae: $\text{DW} = \text{FW} \times \text{plogis}[-0.3081 - 0.2366 \times \ln(A)]$ | | | | | | | | | |
| General: $\text{DW} = \text{FW} \times \text{plogis}[1.4023 + 0.4203 \times \ln(A) - 0.8017 \times \ln(L)]$ | | | | | | | | | |

### 3.3 Cross-dataset comparison and combined area–mass models

Among the four groups shared between datasets, fresh mass slopes showed no detectable differences for Araneae, Myriapoda or Lumbricina (all *P* ≥ 0.163), but were steeper in this study for Hymenoptera (*P* < 0.001) (Figure 4). Dry mass slopes showed no detectable differences for Araneae, Hymenoptera or Myriapoda (all *P* ≥ 0.265), but differed for Lumbricina (*P* = 0.002). Under common-slope models, intercepts differed in all comparisons (*P* < 0.001) except fresh mass in Myriapoda, for which neither slope nor intercept differed (*P* = 0.204 and 0.239, respectively; Figure 4). At the multi-taxon level, including data source as a fixed effect did not improve model fit for either fresh mass (χ*^2^* = 1.15, *P* = 0.285) or dry mass ( ^2^ = 1.89, *P* = 0.170) after accounting for taxon-specific random intercepts and slopes (Figure 5).

**FIGURE 4.**
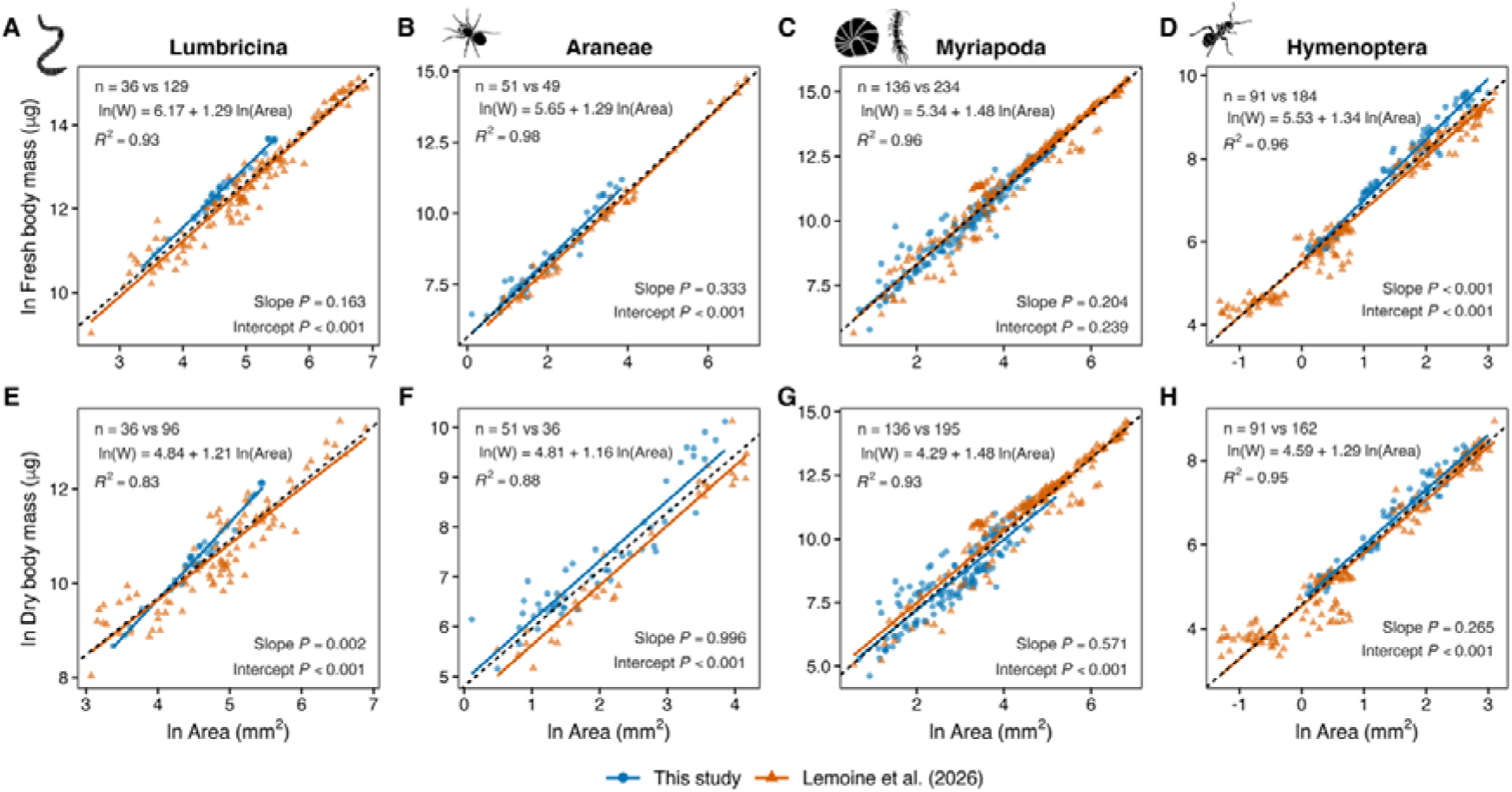
Cross-dataset comparison of area–mass relationships between this study and Lemoine et al. (2026). Fresh mass (**A–D**) and dry mass (**E–H**) are shown for Lumbricina, Araneae, Myriapoda and Hymenoptera. Body area and mass were natural-log transformed. Points and coloured lines represent observations and dataset-specific regressions for this study (blue circles) and Lemoine et al. (2026) (orange triangles); black dashed lines show regressions fitted to the pooled data. Upper-left annotations give sample sizes, pooled regression equations and *R*^2^ values. Lower-right annotations give *P* values for differences in slope and, under common-slope models, intercept between datasets. Soil animal icons were created by S. Meyer.

**FIGURE 5.**
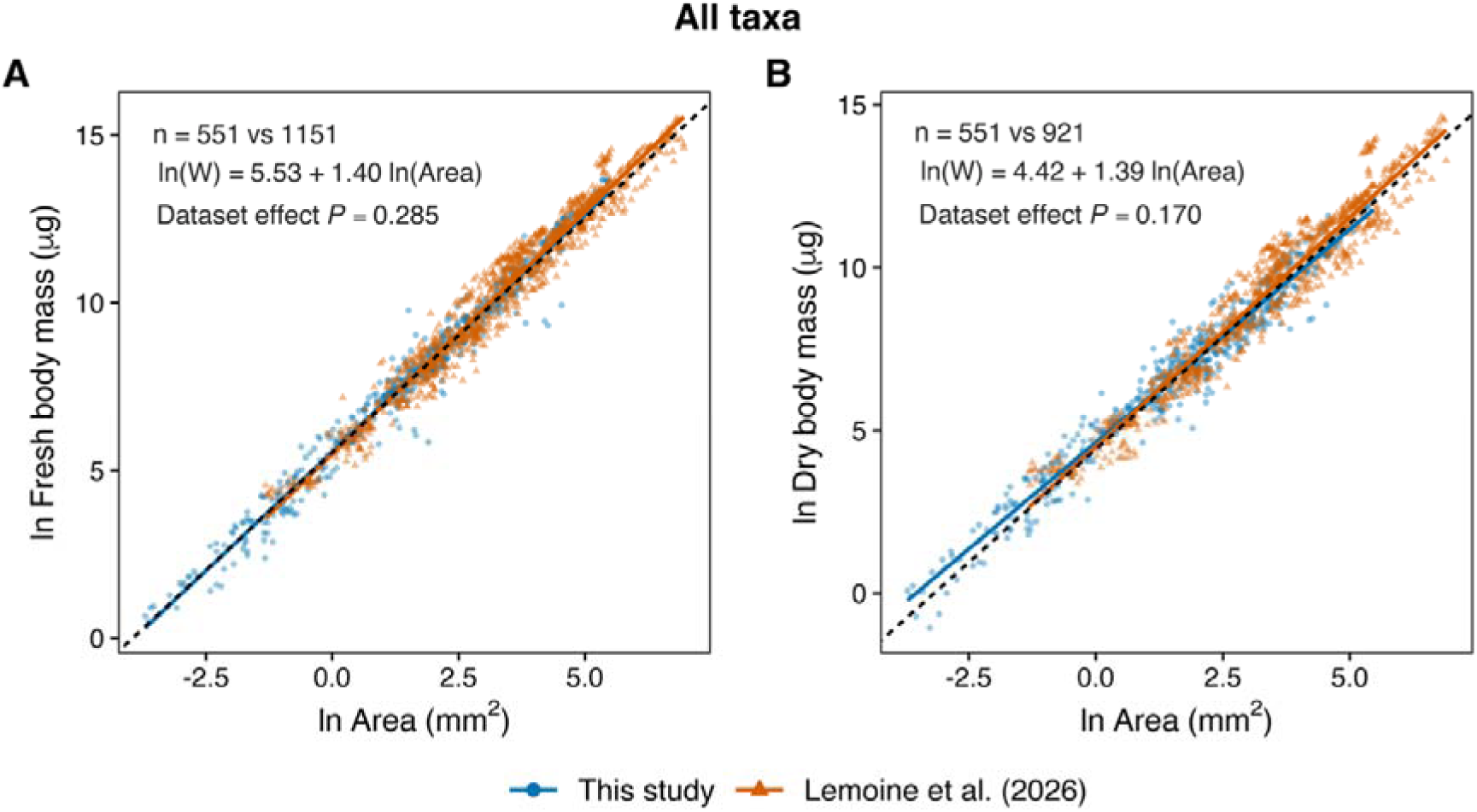
Cross-dataset comparison of area–mass relationships across all taxa. Fresh mass (**A**) and dry mass (**B**) are shown for this study (blue) and Lemoine et al. (2026) (orange), with body area and mass natural-log transformed. Points represent observations, coloured solid lines show dataset-specific regressions, and black dashed lines show population-level predictions from linear mixed-effects models fitted to the combined data with taxon-specific random intercepts and slopes. Panel annotations give sample sizes, population-level area–mass equations and *P* values for the data-source effect.

Pooling both datasets increased sample sizes for shared taxa and expanded taxonomic coverage, yielding combined taxon-specific and general area–mass equations (Table 4). Oribatida and Collembola occurred only in this study, whereas Malacostraca and Gastropoda occurred only in Lemoine et al. (2026). For fresh mass, all taxon-specific models had lower LOOCV MAPE than the general model (16.2–37.3% vs. 40.6%). For dry mass, taxon-specific models also had lower MAPE (21.9–48.9% vs. 55.5%), except for Myriapoda (56.1%). Adjusted *R*^2^ ranged from 0.85 to 0.98 for fresh mass and from 0.83 to 0.97 for dry mass.

**Table 4.**
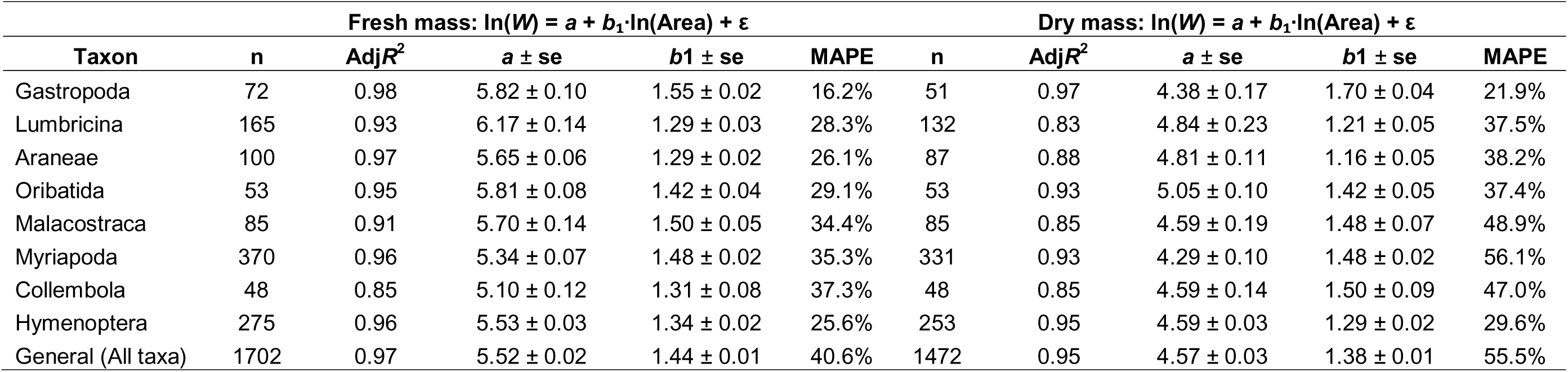
General and taxon-specific area–mass models for fresh mass (left) and dry mass (right) fitted to the combined datasets from this study and Lemoine et al. (2026). All models followed M1: ln(*W*) = *a* + *b*₁ln(*A*) + ε. The general model included all individuals. Shared taxa (Lumbricina, Araneae, Myriapoda and Hymenoptera) were pooled across studies; Oribatida and Collembola occurred only in this study, and Malacostraca and Gastropoda only in Lemoine et al. (2026). *n*, sample size; SE, standard error; adj*R*^2^, adjusted coefficient of determination; MAPE, mean absolute percentage error from leave-one-out cross-validation.

| Fresh mass: $\ln(W) = a + b_1 \cdot \ln(\text{Area}) + \varepsilon$ | | | | | | Dry mass: $\ln(W) = a + b_1 \cdot \ln(\text{Area}) + \varepsilon$ | | | | |
| --- | --- | --- | --- | --- | --- | --- | --- | --- | --- | --- |
| Taxon | n | $\text{Adj}R^2$ | $a \pm \text{se}$ | $b1 \pm \text{se}$ | MAPE | n | $\text{Adj}R^2$ | $a \pm \text{se}$ | $b1 \pm \text{se}$ | MAPE |
| Gastropoda | 72 | 0.98 | $5.82 \pm 0.10$ | $1.55 \pm 0.02$ | 16.2% | 51 | 0.97 | $4.38 \pm 0.17$ | $1.70 \pm 0.04$ | 21.9% |
| Lumbricina | 165 | 0.93 | $6.17 \pm 0.14$ | $1.29 \pm 0.03$ | 28.3% | 132 | 0.83 | $4.84 \pm 0.23$ | $1.21 \pm 0.05$ | 37.5% |
| Araneae | 100 | 0.97 | $5.65 \pm 0.06$ | $1.29 \pm 0.02$ | 26.1% | 87 | 0.88 | $4.81 \pm 0.11$ | $1.16 \pm 0.05$ | 38.2% |
| Oribatida | 53 | 0.95 | $5.81 \pm 0.08$ | $1.42 \pm 0.04$ | 29.1% | 53 | 0.93 | $5.05 \pm 0.10$ | $1.42 \pm 0.05$ | 37.4% |
| Malacostraca | 85 | 0.91 | $5.70 \pm 0.14$ | $1.50 \pm 0.05$ | 34.4% | 85 | 0.85 | $4.59 \pm 0.19$ | $1.48 \pm 0.07$ | 48.9% |
| Myriapoda | 370 | 0.96 | $5.34 \pm 0.07$ | $1.48 \pm 0.02$ | 35.3% | 331 | 0.93 | $4.29 \pm 0.10$ | $1.48 \pm 0.02$ | 56.1% |
| Collembola | 48 | 0.85 | $5.10 \pm 0.12$ | $1.31 \pm 0.08$ | 37.3% | 48 | 0.85 | $4.59 \pm 0.14$ | $1.50 \pm 0.09$ | 47.0% |
| Hymenoptera | 275 | 0.96 | $5.53 \pm 0.03$ | $1.34 \pm 0.02$ | 25.6% | 253 | 0.95 | $4.59 \pm 0.03$ | $1.29 \pm 0.02$ | 29.6% |
| General (All taxa) | 1702 | 0.97 | $5.52 \pm 0.02$ | $1.44 \pm 0.01$ | 40.6% | 1472 | 0.95 | $4.57 \pm 0.03$ | $1.38 \pm 0.01$ | 55.5% |

### 3.4 Community biomass validation

Community biomass predictions were strongly related to measured biomass across all model sets (*R*^2^ = 0.87–0.99, all *P* < 0.001; Figure 6), but prediction error differed substantially among models. Taxon-specific models yielded MAPEs of 16.1–27.3% and slightly overestimated biomass overall (mean ratios = 1.08–1.25). General models showed higher errors, particularly for mesofauna (MAPE = 35.6–42.1%), whereas models fitted to the combined dataset improved mesofauna predictions (MAPE = 9.6% for fresh and 13.9% for dry biomass) but performed poorly for macrofauna, especially dry biomass (MAPE = 52.5%; mean ratio = 0.48).

**FIGURE 6.**
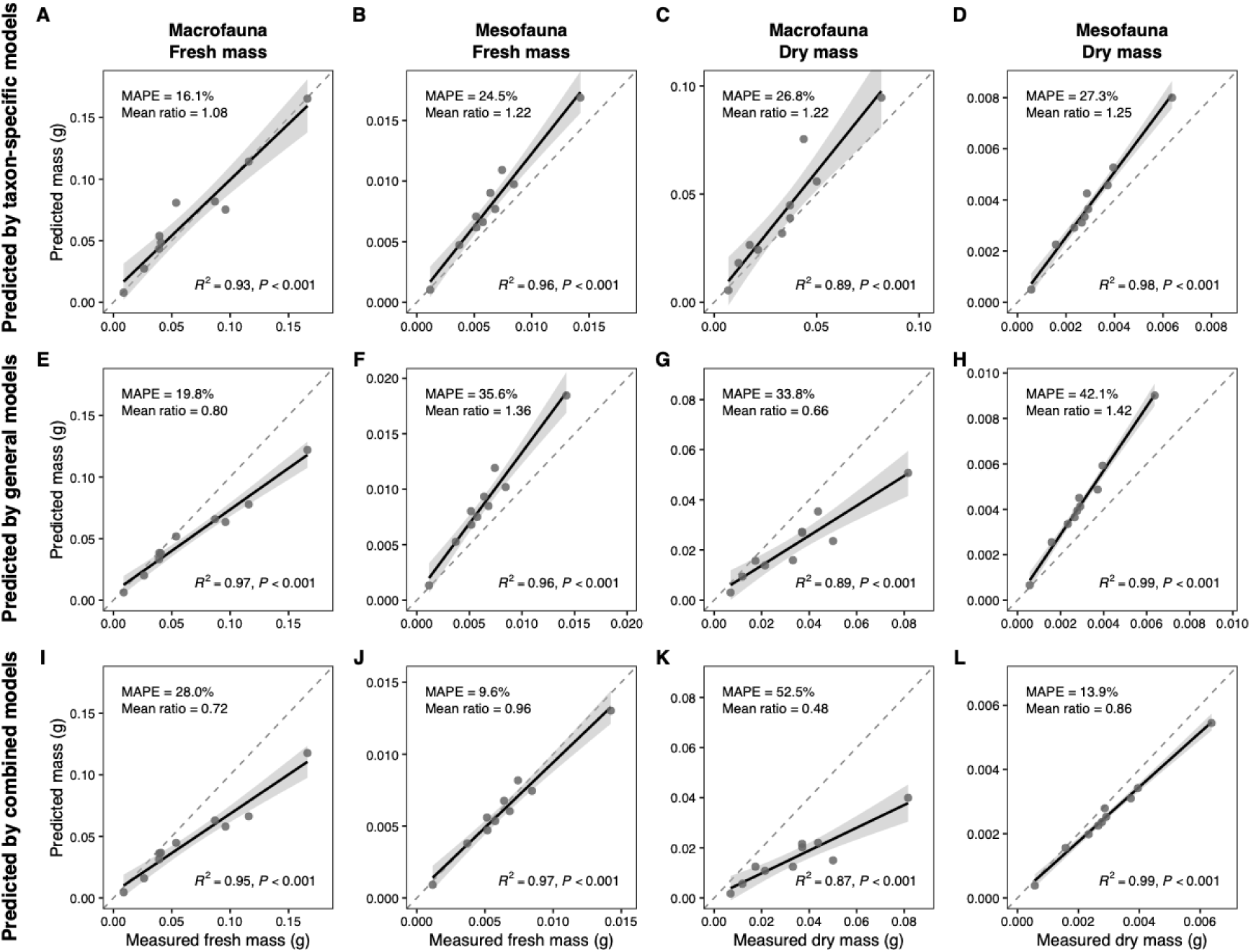
Community-level validation of fresh and dry biomass predictions using taxon-specific, general and combined models. Predicted versus measured biomass is shown for taxon-specific models (**A–D**), the General model (**E–H**) and models fitted to the combined dataset with Lemoine et al. (2026) (**I–L**), for macrofauna and mesofauna fresh and dry biomass (*n* = 10 communities per panel). Solid lines and grey bands show linear regressions with 95% confidence intervals; dashed lines indicate the 1:1 relationship. Panel annotations report *R*^2^, *P*, mean absolute percentage error (MAPE) and the mean predicted-to-measured biomass ratio.

## 4 | DISCUSSION

Our results show that image-derived body area and contour length can be used to estimate fresh and dry mass across morphologically diverse soil invertebrates.

Predictive accuracy varied among taxa, with taxon-specific models generally reducing relative error compared with hierarchical general models, indicating that taxonomic identity provides predictive information beyond body dimensions alone. Cross-dataset comparison with Lemoine et al. (2026) revealed no detectable slope differences for most shared combinations of taxon and mass type and no overall data-source effect after accounting for taxonomic variation, although several taxon-specific differences remained. Pooling the two datasets expanded the model-development dataset and supported area–mass equations across a broader taxonomic range. Finally, community-level validation showed that predicted biomass tracked measured biomass in both macrofauna and mesofauna, although prediction error varied among model sets and faunal groups. Together, these findings support image-based morphometrics as a scalable approach to soil invertebrate biomass estimation while highlighting the continued value of taxon-specific models for predictive accuracy.

### 4.1 Taxon-specific models

Taxon-specific models performed well across morphologically contrasting soil invertebrates, although prediction error varied among taxa. This is particularly relevant for mesofauna, whose biomass has traditionally been estimated from body or tergite length in Collembola and from length-based relationships in Oribatida (Caballero *et al*. 2004; Newton & Proctor 2013; Petersen 1975). Previous image-based area calibrations have focused mainly on macrofauna (Ärje *et al*. 2020; Coulis & Joly 2017; Lemoine *et al*. 2026), making this, to our knowledge, the first direct calibration of projected body area against mass for Collembola and Oribatida.

Prediction error was highest for Collembola but substantially lower for Oribatida (Figure 2), indicating that uncertainty in small-bodied taxa cannot be explained by body size alone. Differences in body architecture and composition are also likely to matter because the same projected area can represent different body volumes and tissue composition, consistent with broader variation in allometric scaling among terrestrial invertebrates (Ehnes *et al*. 2011). Cuticular mineralisation in some Oribatida, for example, illustrates compositional variation not captured by projected area (Norton & Behan-Pelletier 1991). Additional uncertainty may arise from specimen orientation and gut contents, both of which alter the relationship between projected shape and measured mass (Ärje *et al*. 2020; Martin 1986). Taxon-specific equations should therefore be applied within the body-size and imaging conditions represented by their calibration data, with particular caution for the smallest mesofauna, for which reference masses are difficult to measure precisely.

### 4.2 General models

The hierarchical general models captured broad variation in body mass across taxa, with LOOCV *R*^2^ values of 0.97 for fresh mass and 0.96 for dry mass, but retained MAPEs of 27.2% and 22.4%, respectively (Figure 3). This apparent contrast highlights the importance of distinguishing explained variation from predictive error. Because the pooled dataset spans several orders of magnitude in body mass, a high *R*² can arise from correctly capturing broad differences among taxa while appreciable proportional error remains for individual predictions; model performance should therefore not be judged from *R*^2^ alone (Piñeiro *et al*. 2008). The hierarchical structure partly accommodates this heterogeneity by allowing taxon-specific intercepts, and for fresh mass, slopes, while retaining a shared relationship across groups, an important advantage of mixed-effects approaches for heterogeneous ecological data (Bolker *et al*. 2009). Nevertheless, the higher MAPE relative to most taxon-specific models shows that taxonomic differences in mass to morphology scaling are not fully absorbed by a common relationship. This is consistent with large-scale arthropod analyses showing improved mass predictions when taxonomic identity is incorporated into allometric models (Sohlström *et al*. 2018), whereas general equations can perform well within more morphologically constrained groups such as European spiders (Penell *et al*. 2018). In practice, the hierarchical model therefore provides a useful alternative for taxa represented in the model-development dataset but lacking sufficient observations for a separate equation, while taxon-specific models remain preferable when adequate calibration data are available. Its performance for taxa absent from the model-development dataset remains untested and should not be assumed.

### 4.3 Cross-dataset transferability and combined area–mass equations

Comparison with Lemoine et al. (2026) showed that the scaling of mass with projected area was relatively consistent across studies, with slope differences detected only for fresh mass in Hymenoptera and dry mass in Lumbricina. In contrast, intercepts differed in five of the six comparisons with a common slope, suggesting that the scaling relationship was more stable than the absolute mass predicted for a given area (Figure 4). These shifts may reflect both biological and methodological differences. Geographic variation in invertebrate mass–size relationships has been reported previously (Sohlström *et al*. 2018), while specimen processing also differed between studies. Lemoine et al. (2026) scanned freshly collected specimens before long-term ethanol preservation, whereas our specimens were imaged after preservation in 70% ethanol, which may contribute to taxon-dependent changes in body dimensions. At the multi-taxon level, however, data source had no detectable overall effect after accounting for taxon-specific variation, indicating that between-study differences were not uniform across taxa. Pooling both datasets broadened the calibration base, but taxon-specific models still generally showed lower prediction error than the general model, reinforcing the value of retaining taxonomic structure when combining datasets. Transferability remains untested for Collembola and Oribatida, which occurred only in our dataset. These results support combining datasets within taxa to broaden calibration while validating equations independently before applying them beyond the taxonomic or imaging conditions represented in the calibration data.

### 4.4 Validation at the community-biomass scale

Independent validation showed that predicted biomass tracked measured community biomass across all model sets, but absolute prediction error and bias varied markedly among models and faunal groups (Figure 6). Taxon-specific models showed the most consistent performance across macrofauna and mesofauna, whereas the General model produced larger errors for mesofauna. Models fitted to the combined dataset improved mesofauna predictions but underestimated macrofauna biomass, particularly dry biomass. This variation indicates that strong correspondence among communities does not necessarily imply accurate absolute biomass estimates and reinforces the importance of evaluating both goodness of fit and prediction error. These results should be interpreted cautiously because validation was based on only 10 communities per faunal group, limiting inference across broader ranges of community composition and biomass. Preservation may represent an additional source of uncertainty when the equations are transferred among imaging protocols, because ethanol can alter body dimensions in a taxon-dependent manner (Lemoine *et al*. 2026). In our workflow, ethanol was used because it is a standard medium for storing soil invertebrates, and both model-development and validation samples were imaged under the same preservation conditions, making preservation unlikely to explain the observed within-study bias. Despite these limitations, the community validation supports scaling image-derived individual mass estimates to community biomass while retaining individual-level information for biomass-based, trait-based and size-structured analyses of soil-fauna communities.

### 4.5 Limitations and future directions

By establishing area–mass models across major meso- and macrofaunal groups, including the first direct projected-area calibrations for Collembola and Oribatida, and validating predictions at community and cross-dataset scales, our study broadens the taxonomic and empirical basis of image-based biomass estimation for soil fauna. Further generalisation will require broader calibration, because several ecologically important groups, including Enchytraeidae and Coleoptera larvae, were represented by too few individuals for separate taxon-specific models, and the model-development dataset was collected within a single geographic region and sampling season. Cross-dataset comparison with Lemoine et al. (2026) provides partial external validation for several macrofaunal groups, whereas comparable tests are still lacking for Collembola and Oribatida. Broader calibration across regions, populations and environmental contexts will therefore be important, particularly because geographic origin can affect invertebrate mass–size relationships and prediction accuracy (Sohlström *et al*. 2018). A second constraint on scalability is the manual intervention still required for binary-mask generation, separation of touching individuals and taxonomic assignment, although subsequent object detection and extraction of area and contour length are automated using OpenCV. Recent computer-vision approaches provide clear opportunities to reduce this bottleneck: *flatbug* enables automated detection and instance segmentation across diverse terrestrial-arthropod images, while BIODISCOVER and CollembolAI demonstrate automated imaging and classification of arthropod and soil-mesofauna samples (Ärje *et al*. 2020; Svenning *et al*. 2026; Sys *et al*. 2022). Integrating these tools with taxon-specific area–mass equations could enable higher-throughput, taxon-resolved estimates of individual and community biomass, extending automated soil-fauna surveys beyond abundance and taxonomic composition. Such biomass estimates could in turn provide scalable inputs for analyses of metabolic demand and energy flux in soil food webs (Brown *et al*. 2004; Jochum *et al*. 2021; Potapov 2022) and support monitoring of biomass change alongside taxonomic diversity (Eisenhauer *et al*. 2026; Guerra *et al*. 2021).

## ACKNOWLEDGEMENTS

AZ and AP acknowledge support from the Deutsche Forschungsgemeinschaft (DFG, German Research Foundation)—Project number 493345801.

## AUTHOR CONTRIBUTIONS

Anton Potapov and Clement Schneider provided conceptualization. Clement Schneider and Ajuan Zhang developed the statistical methods and wrote the code to implement these methods. Clement Schneider, Ajuan Zhang, Andrey G. Zuev, Cornelia Wiesener, Andreas Kauk, Birgit Balkenhol, Julia Noe, and Sebastian Moll collected and organized the data. Ajuan Zhang analysed the data with advice from Anton Potapov and Clement Schneider. The manuscript was written by Ajuan Zhang with advice from Anton Potapov, Clement Schneider and Jingzhong Lu. All authors contributed critically to drafts of the manuscript and gave final approval for publication.

## DATA AVAILABILITY STATEMENT

Raw data used in the analysis and model code are available from figshare:.

## Notes

### Competing Interest Statement

The authors have declared no competing interest.

## References

Ärje, J., Melvad, C., Jeppesen, M.R., Madsen, S.A., Raitoharju, J., Rasmussen, M.S., et al. (2020). Automatic image-based identification and biomass estimation of invertebrates. Methods Ecol. Evol., 11, 922–931.

Bolker, B.M., Brooks, M.E., Clark, C.J., Geange, S.W., Poulsen, J.R., Stevens, M.H.H., et al. (2009). Generalized linear mixed models: a practical guide for ecology and evolution. Trends Ecol. Evol., 24, 127–135.

Bonfanti, J., Potapov, A.M., Angst, G., Ganault, P., Briones, M.J.I., Calderón-Sanou, I., et al. (2024). Linking effect traits of soil fauna to processes of organic matter transformation. Funct. Ecol., 39(2), 446–461.

Brown, J.H., Gillooly, J.F., Allen, A.P., Savage, V.M. & West, G.B. (2004). TOWARD A METABOLIC THEORY OF ECOLOGY. Ecology, 85, 1771–1789.

Caballero, M., Baquero, E., Ariño, A.H. & Jordana, R. (2004). Indirect biomass estimations in Collembola. Pedobiologia (Jena)., 48, 551–557.

Coulis, M. & Joly, F. (2017). Allometric equations for estimating fresh biomass of five soil macroinvertebrate species from neotropical agroecosystems. Eur. J. Soil Biol., 83, 18–26.

Crowther, T.W., van den Hoogen, J., Wan, J., Mayes, M.A., Keiser, A.D., Mo, L., et al. (2019). The global soil community and its influence on biogeochemistry. Science (80)., 365.

Ehnes, R.B., Rall, B.C. & Brose, U. (2011). Phylogenetic grouping, curvature and metabolic scaling in terrestrial invertebrates. Ecol. Lett., 14, 993–1000.

Eisenhauer, N., Ochoa-Hueso, R., Huang, Y., Barry, K.E., Gebler, A., Guerra, C.A., et al. (2023). Ecosystem consequences of invertebrate decline. Curr. Biol., 33, 4538–4547.e5.

Eisenhauer, N., Sünnemann, M., Pollierer, M.M., Sun, X., Bardgett, R.D., Bartkowski, B., et al. (2026). Soil biodiversity effects on ecosystems. Nat. Rev. Biodivers.

Ellwood, M.D.F. & Foster, W.A. (2004). Doubling the estimate of invertebrate biomass in a rainforest canopy. Nature, 429, 549–551.

Frouz, J. (2018). Effects of soil macro- and mesofauna on litter decomposition and soil organic matter stabilization. Geoderma, 332, 161–172.

Gauzens, B., Barnes, A., Giling, D.P., Hines, J., Jochum, M., Lefcheck, J.S., et al. (2019). fluxweb: An R package to easily estimate energy fluxes in food webs. Methods Ecol. Evol., 10, 270–279.

Guerra, C.A., Bardgett, R.D., Caon, L., Crowther, T.W., Delgado-Baquerizo, M., Montanarella, L., et al. (2021). Tracking, targeting, and conserving soil biodiversity: A monitoring and indicator system can inform policy. Science (80)., 371, 239–241.

Hódar, J.A. (1996). The use of regression equations for estimation of arthropod biomass in ecological studies. Acta Ecologica, 1996, 17, 421, 433.

Höfer, H. & Ott, R. (2009). Estimating Biomass of Neotropical Spiders and Other Arachnids (Araneae, Opiliones, Pseudoscorpiones, Ricinulei) by Mass-Length Regressions. J. Arachnol., 37, 160–169.

Huxley, J.S., Churchill, F.B. & Strauss, R.E. (1993). Problems of relative growth. Johns Hopkins University Press.

Jochum, M., Barnes, A.D., Brose, U., Gauzens, B., Sünnemann, M., Amyntas, A., et al. (2021). For flux’s sake: General considerations for energy-flux calculations in ecological communities. Ecol. Evol., 11, 12948–12969.

van Klink, R., Bowler, D.E., Gongalsky, K.B., Swengel, A.B., Gentile, A. & Chase, J.M. (2020). Meta-analysis reveals declines in terrestrial but increases in freshwater insect abundances. Science (80)., 368, 417–420.

Lemoine, M., Lechopier, T., Marichal, R., Molin, N., & Coulis, M. (2026). Estimating the biomass of soil macrofauna communities using composite images. European Journal of Soil Biology, 130, 103854.

Martin, C.A., Proulx, R. & Magnan, P. (2014). The biogeography of insects’ length– dry mass relationships. Insect Conserv. Divers., 7, 413–419.

Martin, N.A. (1986). Earthworm biomass: Influence of gut content and formaldehyde preservation on live-to-dry weight ratios of three common species of pasture lumbricidae. Soil Biol. Biochem., 18, 245–250.

Newton, J.S. & Proctor, H.C. (2013). A fresh look at weight-estimation models for soil mites (Acari). Int. J. Acarol., 39, 72–85.

Norton, R.A. & Behan-Pelletier, V.M. (1991). Calcium carbonate and calcium oxalate as cuticular hardening agents in oribatid mites (Acari: Oribatida). Can. J. Zool., 69, 1504–1511.

Penell, A., Raub, F. & Höfer, H. (2018). Estimating biomass from body size of European spiders based on regression models. J. Arachnol., 46, 413–419.

Petersen, H. (1975). Estimation of dry weight, fresh weight, and calorific content of various Collembolan species. Pedobiologia (Jena)., 15, 222–243.

Piñeiro, G., Perelman, S., Guerschman, J.P. & Paruelo, J.M. (2008). How to evaluate models: Observed vs. predicted or predicted vs. observed? Ecol. Modell., 216, 316–322.

Potapov, A.M. (2022). Multifunctionality of belowground food webs: resource, size and spatial energy channels. Biol. Rev., 97, 1691–1711.

Potapov, A.M., Sun, X., Barnes, A.D., Briones, M.J., Brown, G.G., Cameron, E.K., et al. (2022). Global monitoring of soil animal communities using a common methodology. Soil Org., 94, 55–68.

Rosenberg, Y., Bar-On, Y.M., Fromm, A., Ostikar, M., Shoshany, A., Giz, O., et al. (2023). The global biomass and number of terrestrial arthropods. Sci. Adv., 9, 1–12.

Schoener, T.W. (1980). Length-Weight Regressions in Tropical and Temperate Forest-Understory Insects. Ann. Entomol. Soc. Am., 73, 106–109.

Seibold, S., Gossner, M.M., Simons, N.K., Blüthgen, N., Müller, J., Ambarlı, D., et al. (2019). Arthropod decline in grasslands and forests is associated with landscape-level drivers. Nature, 574, 671–674.

Sohlström, E.H., Marian, L., Barnes, A.D., Haneda, N.F., Scheu, S., Rall, B.C., et al. (2018). Applying generalized allometric regressions to predict live body mass of tropical and temperate arthropods. Ecol. Evol., 8, 12737–12749.

Svenning, A., Mougeot, G., Alison, J., Chevalier, D., Molina, N.C., Ong, S.Q., et al. (2026). A general method for detection and segmentation of terrestrial arthropods in images. Methods Ecol. Evol., 17, 727–739.

Sys, S., Weißbach, S., Jakob, L., Gerber, S. & Schneider, C. (2022). CollembolAI, a macrophotography and computer vision workflow to digitize and characterize samples of soil invertebrate communities preserved in fluid. Methods Ecol. Evol., 13, 2729–2742.

Winfree, R., Reilly, J.R. & Genung, M.A. (2025). Biodiversity–ecosystem function research must consider abundance and not just diversity. Nat. Rev. Biodivers., 1, 347–348.

Wu, D., Du, E., Eisenhauer, N., Mathieu, J. & Chu, C. (2025). Global engineering effects of soil invertebrates on ecosystem functions. Nature, 640, 120–129.

